# Motor module generalization across balance and walking is reduced after stroke

**DOI:** 10.1101/381939

**Authors:** Jessica L. Allen, Trisha M. Kesar, Lena H. Ting

## Abstract

Here, we examined features of muscle coordination associated with reduced walking performance in chronic stroke survivors. Using motor module (a.k.a. muscle synergy) analysis, we identified differences in the modular control of overground walking and standing reactive balance in stroke survivors compared to age-similar neurotypical controls. In contrast to previous studies that demonstrated reduced motor module number post-stroke, our cohort of stroke survivors did not exhibit a reduction in motor module number compared to controls during either walking or reactive balance. Instead, the pool of motor modules common to walking and reactive balance was smaller, suggesting a reduction in generalizability of motor module function across behaviors. The motor modules common to walking and reactive balance tended to be less variable and more distinct, suggesting more reliable output compared to motor modules specific to one behavior. Indeed, higher levels of motor module generalization was associated with faster walking speeds in stroke survivors. Further, recruitment of a common independent plantarflexor module across both behaviors was associated with faster walking speeds. Our work is the first to show that motor module generalization across walking and balance may help to distinguish important and clinically-relevant differences in walking performance across stroke survivors that would have been overlooked by examining only a single behavior. Finally, as similar relationships between motor module generalization and walking performance have been demonstrated in healthy young adults and individuals with Parkinson’s disease, our work suggests that motor module generalization across walking and balance may be important for well-coordinated walking.

**New and Noteworthy:** Our study is the first to simultaneously examine neuromuscular control of walking and standing reactive balance in stroke survivors. We show that motor module generalization across these behaviors (i.e., recruiting common motor modules) is reduced compared to neurotypical controls, which is associated with slower walking speeds. This is true despite no difference in motor module number between groups within each behavior, suggesting that motor module generalization across walking and balance is important for well-coordinated walking.

## Introduction

More than 50% of stroke survivors are left with mobility impairments that limit their quality of life (Mayo et al. 1999; Miller et al. 2010) and within the first year following a stroke up to 70% of community dwelling stroke survivors experience a fall due to loss of balance (Weerdesteyn et al. 2008). Appropriate muscle coordination is required for well-coordinated walking and maintaining balance, particularly in response to external perturbations such as slips and trips. However, muscle coordination is often impaired after stroke for both gait (Knutsson and Richards 1979; Shiavi et al. 1987; Clark et al. 2010) and balance control (Di Fabio et al. 1986; Kirker et al. 2000; Marigold and Eng 2006; de Kam et al. 2018). Motor module (a.k.a. muscle synergy) analysis has proven useful in providing insight into changes in muscle coordination associated with reduced walking performance in neurological populations such as stroke (Ivanenko et al. 2013; Ting et al. 2015; Seamon et al. 2018). Motor modules are groups of coactive muscles that are flexibly recruited over time to transform movement goals into biomechanical outputs. To date, motor module analysis for lower limb muscle coordination post-stroke has primarily focused on the number of motor modules recruited during gait behaviors. We recently utilized novel metrics of motor module consistency, distinctness, and generalization to examine other features of muscle coordination and identified differences related to gait and balance performance in both healthy adults (Sawers et al. 2015) and individuals with Parkinson’s disease (Allen et al. 2017). However, it remains unclear how these modular features of muscle coordination are affected after stroke. Identifying features of muscle coordination associated with impaired gait and balance performance post-stroke through motor module analysis may provide important insight into neuromuscular mechanisms underlying impaired motor performance.

Compared to neurotypical controls, stroke survivors often recruit a smaller number of motor modules during walking, i.e., reduced neuromuscular complexity (Clark et al. 2010). This reduction in motor module number is due to merging of different healthy modules, assumed to reflect a lack of independent drive to motor modules that perform different functions. Motor module merging post-stroke is consistent with the clinical definition of muscle synergies, in which abnormal coupling of muscles across the limb are observed (Knutsson and Richards 1979; Shiavi et al. 1987). Further, the reduction in motor module number post-stroke is associated with impaired gait and balance function (Bowden et al. 2010; Clark et al. 2010; Barroso et al. 2017) and limits the ability to perform more complex locomotor tasks (e.g., changing speed, cadence, step length, and step height, Routson et al. 2014). Motor module number is also better correlated with gait and balance function than are lower limb Fugl-Meyer assessments typically used to measure the severity of motor impairment (Bowden et al. 2010). Several studies have also shown that improved walking performance after rehabilitation is associated with increases in motor module number (Routson et al. 2013; Ferrante et al. 2016).

However, the ability to recruit a given number of motor modules does not directly translate to a specific level of motor performance. Although recruiting fewer motor modules is associated with slower walking speeds post-stroke, differences in speed still remain in individuals who recruit an identical number of modules (Clark et al. 2010). Similarly, improved walking performance after post-stroke rehabilitation can occur without increasing motor module complexity (Routson et al. 2013; Hashiguchi et al. 2016). To aid in distinguishing important and clinically relevant impairments in motor performance, we recently introduced novel motor module metrics of consistency and distinctness (Sawers et al. 2015; Allen et al. 2017). These novel metrics reflect variations in muscle coordination across different repetitions of the same behavior (e.g., from gait cycle to gait cycle for walking). We posit that greater motor module consistency and distinctness reflects greater stability of motor output across repetitions of a behavior (consistency) organized around producing more well-defined biomechanical output (distinctness), leading to higher levels of motor performance. Indeed, we recently observed greater motor module consistency and distinctness during a balance-challenging walking behavior among expert professional ballet dancers compared with novice nondancers (Sawers et al. 2015). Similarly, we found that improved gait and balance performance after rehabilitation in individuals with Parkinson’s disease was associated not with increased module complexity but increased consistency and distinctness (Allen et al. 2017). Although it is well-established that stroke survivors walk with increased step-to-step variability (e.g., spatiotemporal variability, Balasubramanian et al. 2009), whether reduced motor performance is stroke survivors is accompanied by reduction in motor module consistency and distinctness is unknown.

Maintaining balance is critical for walking, especially in the presence of external disturbances, yet little is known about motor modules recruited for balance post-stroke and how they compare to modules recruited during walking. Recent evidence suggests that generalization of motor modules across walking and balance behaviors, i.e., recruiting a common set of motor modules, may be an important feature of muscle coordination underlying differences in walking performance. In healthy, young adults, many of the motor modules recruited during walking are also recruited to control balance in response to external perturbations (Chvatal and Ting 2012, 2013; Oliveira et al. 2012). Further, higher levels of motor module generalization across walking and balance behaviors is associated with better motor performance. Long-term training over many years in professional ballet dancers leads to better motor performance on a balance-challenging beam-walking behavior compared to nondancers, which is associated with recruiting more common motor modules across gait and balance movement behaviors (Sawers et al. 2015). On the other end of the motor expertise-impairment spectrum, individuals with Parkinson’s disease exhibit lower levels of motor module generalization across gait and balance behaviors compared to healthy adults, and improvements in motor performance after rehabilitation are associated with increases in motor module generalization (Allen et al. 2017). Whether motor module generalization is reduced in stroke survivors whose ability to selectively recruit appropriate patterns of muscle coordination is impaired (e.g., Clark et al. 2010; Knutsson and Richards 1979; Shiavi, Bugle, and Limbird 198) remains unknown.

In the present study we analyzed electromyography (EMG) data from muscles spanning the hip, knee, and ankle during overground walking and multidirectional perturbations to standing to examine how the modular control of walking and balance is affected in stroke survivors. We hypothesized that having a stroke impairs the ability to selectively and consistently recruit patterns of neuromuscular control appropriate for a given movement behavior. Based on this hypothesis, we predicted that stroke survivors (a) recruit fewer motor modules in walking and in balance that have (b) less consistency and distinctness in their structure, such that (c) fewer common motor modules are recruited across walking and balance behaviors.

## Methods

### Subjects

Nine individuals post-stroke (3 male, 57.2±12.7 years, 85.5±24.4 kg, 6 right-sided hemiparesis, 46.3±23.1 months post-stroke, Fugl-Meyer lower extremity 23.7±3.7) and eight sex-, age-, and weight-similar neurotypical controls (3 male, 62.0±6.6 years, 76.4±19.1 lbs.) participated in the current study (Table 1). All participants provided written informed consent prior to participating according to protocols approved by the institutional review boards at both Emory University and Georgia Institute of Technology.

**Table 1:** Subject demographics

| Stroke Group |  |  |  |  |  |  | Control Group |  |  |  |
| --- | --- | --- | --- | --- | --- | --- | --- | --- | --- | --- |
| Subject | Sex | Age | Mass | Affected | Months | Stroke | Subject | Sex | Age | Mass |
|  |  | (yr) | (kg) | Side | post-stroke | type |  |  | (yr) | (kg) |
| S1 | F | 61 | 44.0 | R | 73 | H | C1 | F | 63 | 49.8 |
| S2 | M | 70 | 74.0 | R | 60 | I | C2 | F | 68 | 62.4 |
| S3 | M | 53 | 98.6 | L | 35 | I | C3 | F | 60 | 86.0 |
| S4 | F | 65 | 97.8 | L | 47 | I | C4 | M | 56 | 113.1 |
| S5 | F | 63 | 85.9 | R | 48 | I | C5 | F | 67 | 81.4 |
| S6 | F | 43 | 133.5 | L | 11 | I | C6 | M | 57 | 81.0 |
| S7 | F | 67 | 72.7 | L | 50 | I | C7 | M | 72 | 73.6 |
| S8 | F | 31 | 88.7 | R | 15 | H | C8 | F | 53 | 64.3 |
| S9 | M | 62 | 74.2 | R | 78 |  |  |  |  |  |

Stroke type: H = hemorrhagic, I = ischemic.

Inclusion/exclusion criteria for individual’s post stroke were: <u>Inclusion</u>: (1) chronic stroke (>6 months post-stroke), (2) first (single) lesion, (3) lower-limb Fugl-meyer >12 and <34, (4) ambulatory with or without an assistive device, and (5) ability to stand unassisted for at least 15 minutes. <u>Exclusion</u>: (1) inability to communicate with investigators, (2) lower extremity joint pain, contractures, major sensory deficits, cardiovascular or respiratory symptoms contra-indicative of walking, (3) any other significant non-stroke-related impairment affecting balance, walking or cognition.

Inclusion/exclusion criteria for neurotypical controls included: <u>Inclusion</u>: (1) ambulatory with or without an assistive device, (2) ability to stand unassisted for at least 15 minutes, and (3) age ≥ 18 years. <u>Exclusion</u>: inability to communicate with researchers, (2) lower extremity joint pain, contractures, major sensory deficits, cardiovascular or respiratory symptoms contra-indicative of walking, (3) history or evidence of orthopedic, muscular, or physical disability, (4) taking current medications that may affect balance, (5) history or evidence of vestibular, auditory, or proprioceptive impairment, (6) history or indication of orthostatic hypotension, (7) history of any neurological disease or insult.

### Experimental protocol

We recorded postural responses to ramp-and-hold translations of the support surface during standing while subjects stood on an instrumented platform that translated in 12 equally-spaced directions in the horizontal plane (see **Fig. 1B**). Subjects were instructed to maintain balance without stepping. Three trials in each direction were collected in random order. All subjects were exposed to the same level of perturbation (displacement 7.5cm, velocity 15cm/s, acceleration 0.1g). This perturbation level was such that all subjects could maintain balance on a majority of trials such that few corrective steps were observed. Stance width was self-selected and enforced to be the same across all trials.

**Figure 1:**
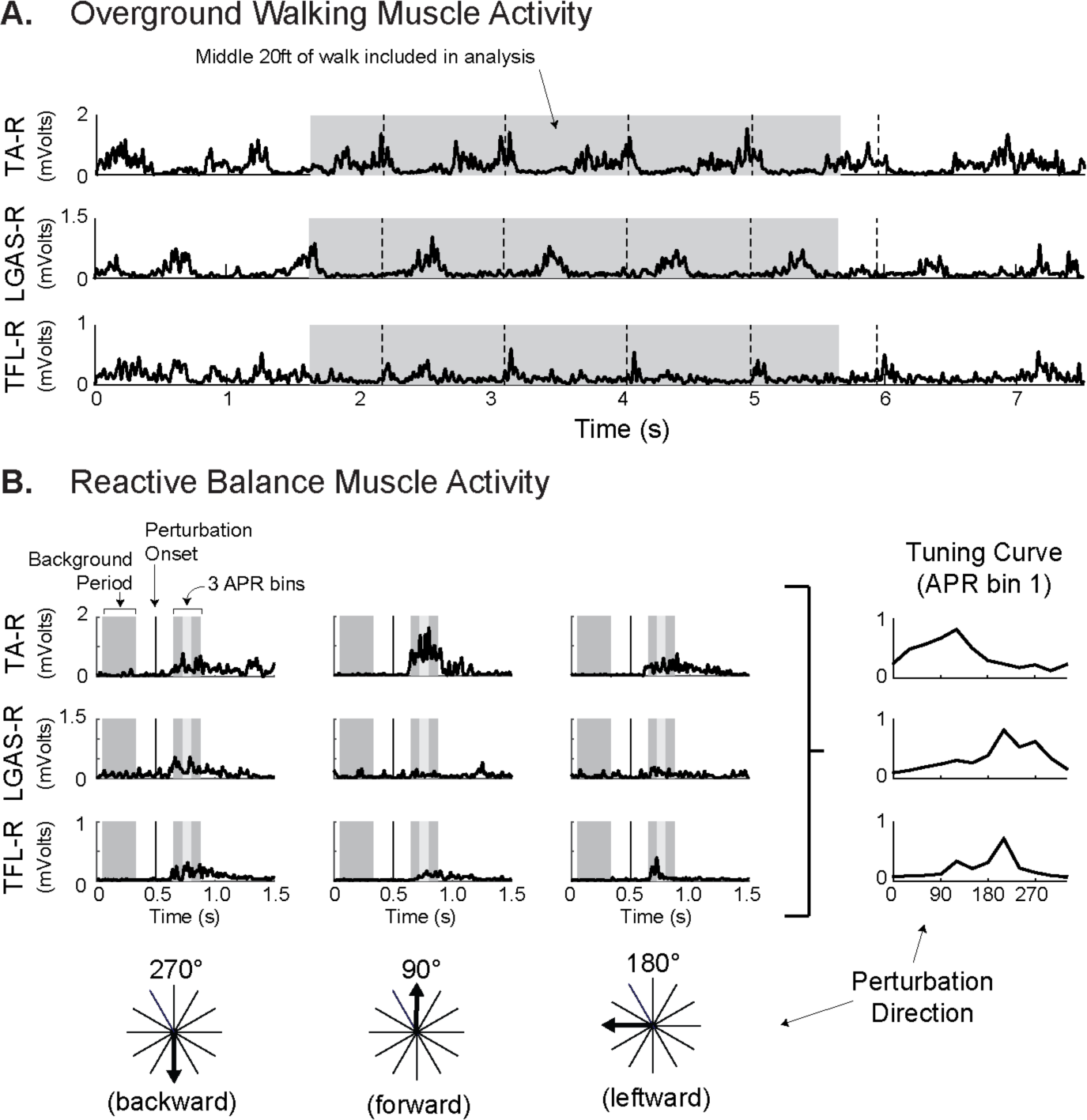
Example processed EMG from select muscles during overground walking (A) and reactive balance (B). A: muscle activity for walking was recorded while participants walked overground at their self-selected speed for at least 3 trials over a 25 ft walkway. Dashed lines represent right heel strikes. For each trial, only data from the middle 20 ft of the 25ft walkway were analyzed to avoid the effects of gait initiation and termination, represented by the shaded region. Data from all trials for a subject were concatenated before motor module extraction to form an *m × t* data matrix, where *m* is the number of muscles and *t* the number of time points across all trials. B: muscle activity for reactive balance was assessed through ramp-and-hold perturbations in 12 evenly spaced directions. Left: responses to backward, forward, and leftward perturbations are illustrated. EMG responses occurred ~150 ms after perturbation onset (denoted by vertical lines). Mean EMG activity was calculated during a background period before the perturbation and during three 75-ms time bins during the automatic postural response (APR, shaded regions). Right: tuning curves of mean muscle activity from perturbation responses as a function of perturbation directions for the first APR bin. Before motor module extraction, the tuning curves were assembled to form an *m × t* data matrix, where *m* is the number of muscles and *t* the number of data points (3 trials × 12 directions ×4 time bins = 144).

Each subject also walked over-ground at self-selected walking speed over a ~25ft distance. Subjects were instructed to walk as they would normally while keeping their head up and looking straight ahead. Walking speed for each trial was defined as the average velocity of the C7 marker in the middle 20 ft of the walkway and averaged across trials to serve as a measure of walking performance.

Surface electromyography was recorded from 12 muscles per leg: gluteus maximus (GMAX), gluteus medius (GMED), tensor fascia lata (TFL), adductor magnus (ADMG), biceps femoris long-head (BFLH), rectus femoris (RFEM), vastus lateralis (VLAT), medial gastrocnemius (MGAS), lateral gastrocnemius (LGAS), soleus (SOL), peroneus longus (PERO), and tibialis anterior (TA). All EMG data were collected at 1200 Hz except for in subjects S1-S4, in which EMG was collected at 1080 Hz. Three-dimensional kinematics were measured at 120 Hz with an eight-camera (subjects S1-S4) or ten-camera (all other subjects) Vicon motion capture system and a custom 25-marker set that included head-arms-trunk, thigh, shank, and foot segments.

### EMG data processing

EMG data were high-pass filtered at 35 Hz, demeaned, rectified, and low-pass filtered at 40 Hz with custom MATLAB routines. Subject specific EMG data matrices for each leg and condition (i.e., walking and reactive balance) were created as follows.

For reactive balance, EMG data were analyzed during four different time bins: one before the perturbation and three during the automatic postural response (APR; **Fig. 1B**) (Chvatal and Ting 2013). Specifically, mean muscle activity was calculated during a 120-ms background period that ended 170-ms before the perturbation and during each of three 75-ms bins beginning either 150-ms after perturbation onset. Mean muscle activity values for each muscle during each bin for each trial were assembled to form an *m×t* data matrix, where *m* is the number of muscles (12) and *t* the number of data points (3 trials × 12 directions × 4 time bins = 144).

For walking, at least 10 gait cycles were analyzed per subject to ensure adequate capture of step-to-step variability in muscle recruitment (Oliveira et al. 2014). For consistency with reactive balance processing, EMG data for walking were averaged over 75-ms bins. Only data from the middle 20 ft of the 25ft walkway were analyzed to avoid the effects of gait initiation and termination (**Fig. 1A**). Trials were concatenated end to end to form an *m×t* data matrix. The number of conditions, *t* (trials × time bins), varied across subjects. The minimum size of *t* was 149, and there was no significant difference in the size of *t* between groups (388.1±139.8 for stroke subjects, 341.6±81.8 for controls; t(15)=0.822, p=0.424).

The assembled EMG data matrices for each condition were normalized to the maximum activation observed during walking at self-selected speed.

### Motor module analysis

Four sets of motor modules were identified for each subject (i.e., 2 legs x 2 conditions [walking and reactive balance]). Motor modules were identified by applying a non-negative matrix factorization algorithm on the EMG data matrices (NNMF, Lee and Seung 1999), such that EMG=W×C. W is an *m×n* matrix with *n* motor modules and C is an *n×t* matrix of motor module activation coefficients. To ensure equal weighting of each muscle during the extraction process, each row in the EMG data matrices (i.e., each muscle) was scaled to unit variance before motor module extraction and rescaled to original units afterwards (Torres-Oviedo and Ting 2007).

The number of motor modules, *n*, per condition was chosen as described in Allen et al. 2017. Briefly, 1-12 motor modules (W) were extracted from each EMG data matrix. The goodness of fit between actual and reconstructed EMG was evaluated with variability accounted for (VAF), defined as 100 × squared uncentered Pearson’s correlation coefficient (Zar 1999). The number of motor modules was chosen such that the lower bound of the 95% confidence interval on VAF exceeded 90% (Cheung et al. 2009; Hayes et al. 2014; Allen et al. 2017). Confidence intervals were calculated using a bootstrapping procedure (250 samples with replacement).

#### Motor module structure was analyzed using the following primary outcome metrics

##### Motor module number (n_walk_, n_balance_)

Motor module complexity was defined as the number of motor modules independently extracted from the EMG data matrices for walking and reactive balance. To test our prediction that individuals post-stroke would exhibit reduced complexity on the paretic leg, we compared the number of motor modules independently extracted in each leg (control, nonparetic, paretic) using separate Kruskall-Wallis tests for walking and reactive balance.

##### Motor module variability (R95_walk_, R95_balance_)

Motor module variability was defined as the variability of motor module structure across different movement observations. This analysis quantifies the variability of motor module spatial structure (W) across different subsets of the EMG data set with a multistep process (similar to Allen et al. 2017). First, each EMG matrix was resampled 100 times in which 80% of the data was randomly sampled without replacement. From each resampled matrix, a new set of motor modules was extracted, where the number of motor modules, *n*, was identical to the number previously identified from the entire data set. Because each extraction does not extract modules in the same order, a k-means algorithm was used to cluster similar modules across the 100 resampled extractions. The initial seed for the k-means algorithm was the motor modules extracted using all the data. The variability of each motor module was then quantified as the radius of the n-sphere (in 12D space) that encompassed the all cluster points (e.g., the 100 different re-sampled motor modules) in that module to 95% confidence, which was then averaged across all modules within a task. For a 2D representation, see **Fig. 3A**. To test our prediction that individuals post-stroke would exhibit increased motor module variability on the paretic leg, we compared motor module variability in each leg (control, nonparetic, paretic) using separate one-way ANOVAs for walking and reactive balance.

**Figure 3:**
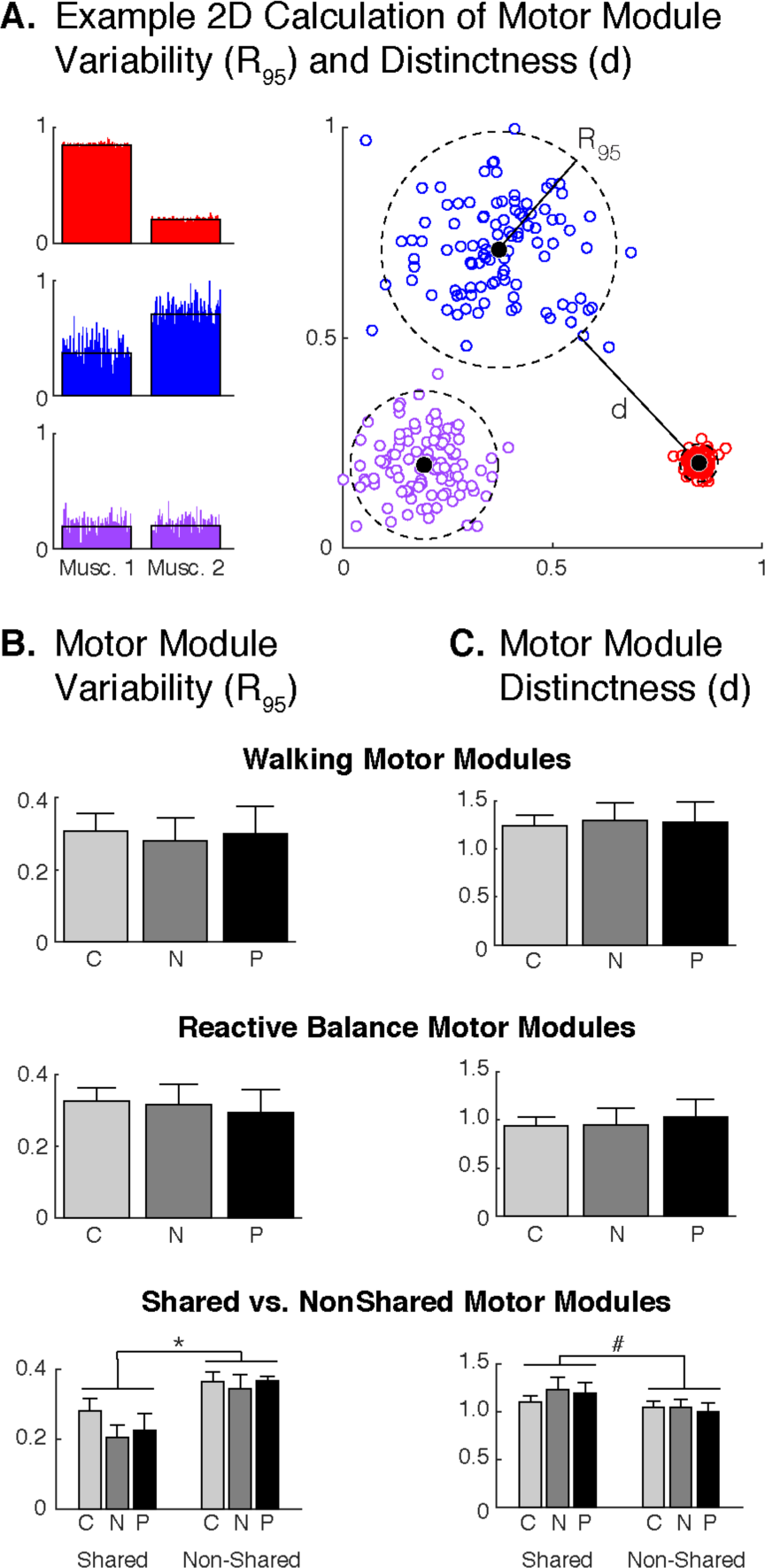
Motor module variability and distinctness. A: A two-dimensional (i.e. 2 muscles) example of motor module variability and distinctness calculation. *Left:* colored bars for each muscle weighting represent the contribution of muscles 1 and 2 within a module over each of the 100 different resampled module extractions. Black bars indicate the mean across all resampled extractions. *Right*: each point in a cluster represents 1 of the 100 resampled motor modules as depicted on *left*. B,C: Motor module variability and distinctness did not differ between control (n=16, light gray), nonparetic (n=9, dark gray), and paretic legs (n=9, black) in either walking or reactive balance. However, in all legs the motor modules that were shared across the two behaviors exhibited less variability and were more distinct than those that were recruited in only one of the behaviors. * and # denote p < 0.05 and p < 0.1, respectively.

##### Motor module distinctness (d_walk_, d_balance_)

Motor module distinctness was defined as the mean distance between the R95 n-spheres of each module in 12D space, where the more distinct the motor modules are for a task the greater the distance. For 2D representative example, see **Fig. 3A**. To test our prediction that individuals post-stroke would exhibit decreased motor module distinctness on the paretic leg, we compared the number of motor modules independently extracted in each leg (control, nonparetic, paretic) using separate one-way ANOVAs for walking and reactive balance.

##### Motor module generalizability (%n_shared_)

Motor module generalizability was defined as the percentage of motor modules that were similar between reactive balance and walking, %n_shared_. First, the number of shared motor modules across walking and reactive balance (*n_shared_*) was identified with Pearson’s correlation coefficients (Chvatal and Ting 2013; Allen et al. 2017). A pair of motor modules were considered “shared” if r≥0.708, which corresponds to the critical value of r^2^ for 12 muscles at p=0.01. The amount of shared motor module was expressed as a percentage to account for the fact that each subject recruited a different number of total motor modules. The percentage of shared motor modules was calculated as 100 × [*n*_shared_/(*n*_walk_+*n*_balance_–*n*_shared_)]. To test our prediction that individual’s post-stroke would have a reduced pool of common motor modules between walking and reactive balance, we compared the percentage of shared motor modules across walking and reactive balance in each leg (control, nonparetic, paretic) using a one-way ANOVA. All statistics were performed in SPSS (version 25; IBM SPSS, Chicago, IL) with α=0.05.

#### The following secondary analyses were also performed

As a secondary analysis on motor module variability and distinctness, we also examined whether variability and distinctness differed between shared vs. non-shared modules using a separate two-way ANOVA with ‘leg’ and ‘shared’ as factors. The leg factor had three levels (Control, Paretic, Nonparetic) and the shared factor had two levels (Shared, NonShared). In this analysis, the variability and distinctness of each individual module was assessed (instead of averaged within a subject) and split into two groups (Shared and Nonshared). Leg was included as a factor to check that a similar leg-effect was identified as in the one-way ANOVA examining the effect of leg as reported above.

To investigate the relationship between generalizability and motor performance in individual post-stroke, several secondary analyses were performed. First, Pearson’s correlation analyses were performed between walking speed and motor module generalization in the paretic leg. We then examined the structure of the motor modules shared between walking and reactive balance. To facilitate comparison of modules between legs, motor modules from walking that were also recruited in reactive balance were pooled across legs and grouped with a hierarchical cluster analysis (MATLAB statistics-toolbox functions pdist [Minkowski option; P = 3], linkage [ward option], and cluster). The number of unique shared modules across legs was determined by identifying the minimum number of clusters that partitioned motor modules such that no cluster contained more than per leg (Cheung 2005; Sawers et al. 2017). Finally, to determine whether the recruitment of any of these shared modules was associated with walking speed, a stepwise linear regression model was created in MATLAB. Six initial regressors were included in the model, corresponding to presence of module 1, module 2, and module 3 in each of the paretic and nonparetic legs. Presence or absence of a shared module was coded as 1 or 0, respectively. Regressors were added or removed based on the p-value of the F-statistic less than or greater than 0.05 and 0.1, respectively.

## Results

Motor module number (**Fig. 2B**) was not significantly different between control, paretic, and nonparetic legs for either walking (p=0.801) or reactive balance (p=0.486). The median number of motor modules recruited for walking was 4 in controls (range: 2-5), 4 in the nonparetic leg (range: 3-5), and 4 in the paretic leg (3-5). The median number of motor modules recruited for reactive balance was 3 in controls (range: 2-4), 3 in the nonparetic leg (range: 3-4), and 4 in the paretic leg (2-5).

**Figure 2:**
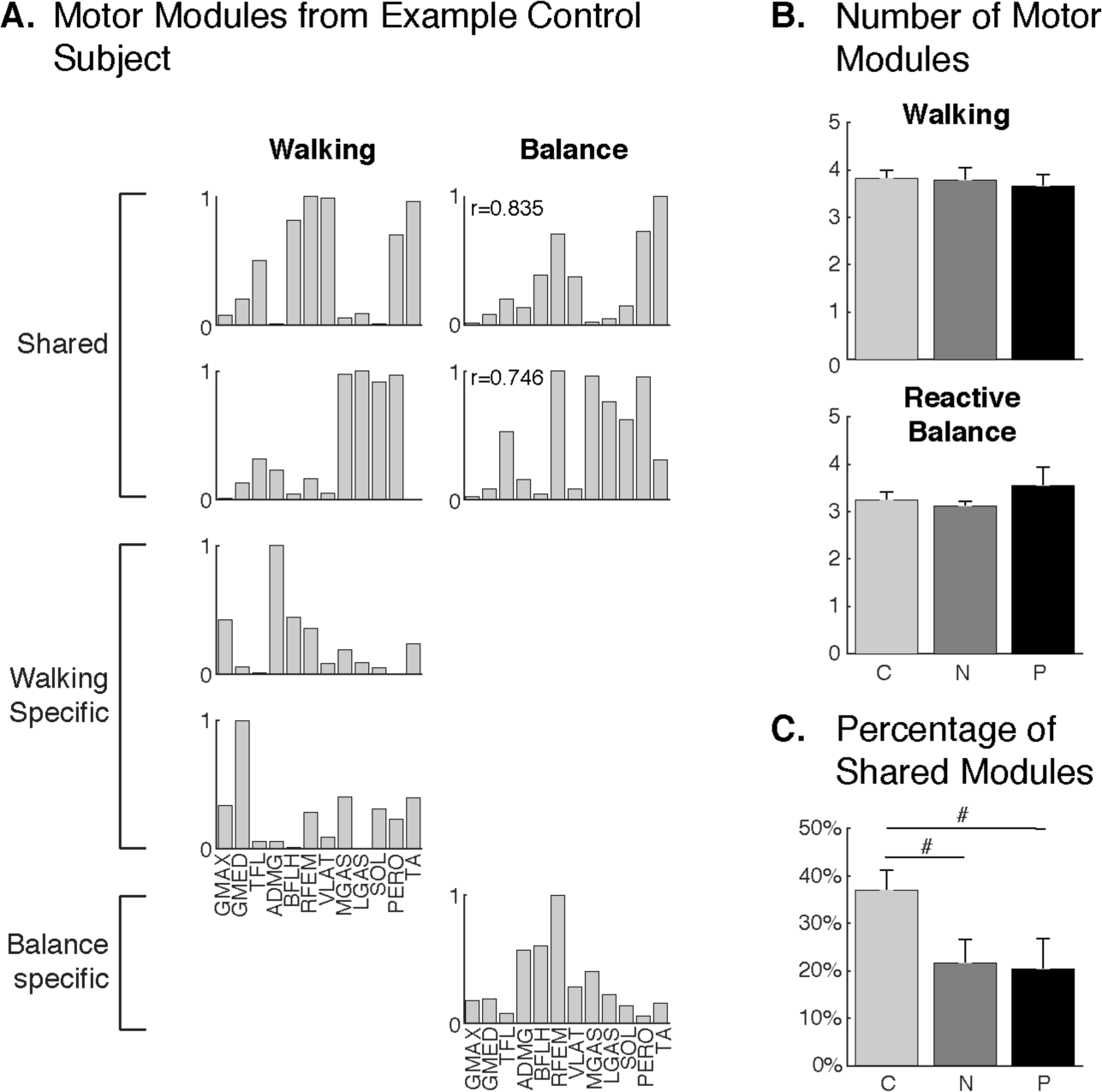
Motor module number and generalization across walking and reactive balance. A: representative motor modules from a control subject during walking (left) and reactive balance (right). Motor modules were extracted from each behavior independently and identified as shared across behaviors if r > 0.708. B: The number of motor modules recruited during overground walking (top) and reactive balance (bottom) did not differ between control (n=16, light gray), nonparetic (n=9, dark gray), and paretic (n=9, black) legs. C: The percentage of shared motor modules was decreased in both the nonparetic and paretic legs compared to control legs. Sharing across behaviors quantified as % of total number of unique motor modules (i.e., 40% of the motor modules, or 2 of 5, were shared across behaviors in the representative subject in A). * and # denote p < 0.05 and p < 0.1, respectively.

We found that motor module variability (**Fig. 3B**) did not differ between control, paretic, or nonparetic legs in either walking (F(2,31)=0.054, p=0.948) or reactive balance (F(2,31)=0.116, p=0.891). Variability for walking across control legs was 0.31±0.20 (range 0.09-0.66), across paretic legs was 0.30±0.23 (range 0.06-0.77), and across nonparetic legs was 0.28±0.20 (range 0.06-0.66). Variability for reactive balance across control legs was 0.33±0.1 (range 0.12-0.60), across paretic legs was 0.29±0.19 (range 0.08-0.63), and across nonparetic legs was 0.32±0.16 (range 0.15-0.63).

Similarly, motor module distinctness (**Fig. 3C**) did not differ between control, paretic, or nonparetic legs in either walking (F(2,31)=0.039, p=0.962) or reactive balance (F(2,31)=0.115, p=0.892). Distinctness for walking across control legs was 1.24±0.45 (range 0.35-1.83), across paretic legs was 1.28±0.63 (range 0.04-1.95), and across nonparetic legs was 1.30±0.56 (range 0.40-1.95). Distinctness for reactive balance across control legs was 0.94±0.39 (range 0.06-1.50), across paretic legs was 1.03±0.57 (range 0.03-1.56), and across nonparetic legs was 0.95±0.52 (range 0.00-1.46).

In contrast, we found significant differences in motor module generalizability (e.g., the percentage of motor modules that were shared between reactive balance and walking) between groups (F(2,31)=3.689, p=0.037; **Fig. 2C**). Motor module generalizability was 37.0±17.0% across control legs (range 12.5-75%), 20.3±19.7% across paretic legs (range 0-50%), and 21.7±14.8% across nonparetic legs (range 0-50%). Between-leg comparisons revealed a trend for reduced motor module generalizability compared to controls in both the paretic (p=0.066) and nonparetic legs (p=0.098), but not between the paretic and nonparetic legs (p=0.984).

Our secondary analyses on motor module variability and distinctness revealed that motor modules common to walking and reactive balance tended to be less variable and more distinct than those that were not common across behaviors. For motor module variability (**Fig. 3B**, bottom panel), we found a significant main effect of shared (F(1,235)=12.433, p=0.001), no effect of leg (F(2,235)=0.728, p=0.484), and no interaction effect (F(2,235)=0.414, p=0.662). Motor module variability was lower in shared vs nonshared modules with a medium effect size (ES=0.45). A lower value of variability means that motor module structure was more consistently recruited from step-to-step. For motor module distinctness (**Fig. 3C**, bottom panel), there was a trend for a main effect of shared (F(1,238)=3.740 p=0.054), and no effect of leg (F(2,238)=0.070, p=0.783) or shared*leg interaction (F(2,238)=0.451, p=0.638). Motor module distinctness was higher in shared vs. nonshared modules, although the effect size was small (ES = 0.22).

Our secondary analyses on motor module generalizability revealed an association with walking performance. We found a moderate positive relationship between motor module generalizability in the paretic leg and walking speed (r=0.46), such that recruiting more common motor modules across walking and reactive balance was associated with walking at faster speeds. A total of three unique shared motor modules were identified across participants (**Fig. 4**), only one of which predicted walking speed. Shared module one primarily consisted of the ankle plantarflexors and was recruited in 13 of 16 control legs, 8 of 9 nonparetic legs, and 4 of 9 paretic legs. Shared module two consisted primarily of the ankle dorsiflexors with low level recruitment of more proximal knee and hip muscles. This module was recruited in 8 of 16 control legs, 0 of 9 nonparetic legs, and 4 of 9 paretic legs. The third shared module consisted of hip, knee, and ankle muscles and was recruited in 7 of 16 control legs, 2 of 9 nonparetic legs, and 2 of 9 paretic legs. Only the presence of shared module 1 (i.e., the plantarflexor module) in the paretic leg was identified as a significant predictor of walking speed (Table 2).

**Table 2:** Regression model relating presence of shared motor modules to walking speed in individuals post-stroke.

|  | Paretic Leg |  | Nonparetic Leg |  |
| --- | --- | --- | --- | --- |
|  | Estimate | p | Estimate | p |
| Module 1 | **0.53** | **0.029** | -0.28 | 0.42 |
| Module 2 | 0.01 | 0.98 | 0.04 | 0.88 |
| Module 3 | -0.19 | 0.38 | 0.00 | 1.0 |

**Figure 4:**
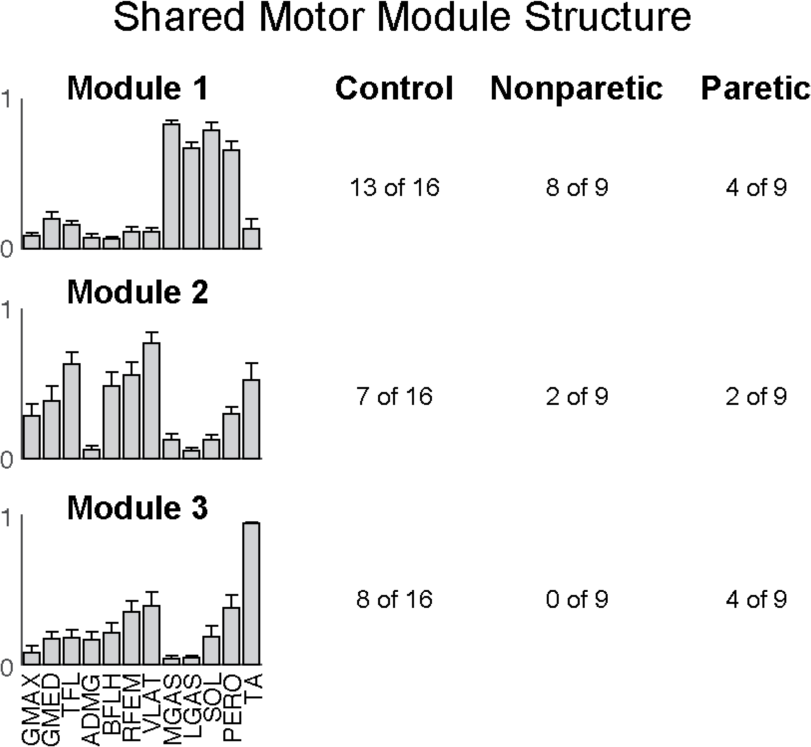
Structure of motor modules shared across walking and reactive balance. Three unique motor modules were identified across all legs as shared across behaviors. The numbers to the right indicate in how many legs per group (control, nonparetic, paretic) each of these motor modules were present.

## Discussion

Here, we show that examining muscle recruitment across movement behaviors with different biomechanics and neural control may reveal important insights into changes in neuromuscular control underlying motor performance that might otherwise be overlooked. Our study is the first to simultaneously examine the modular control of walking and standing reactive balance in stroke survivors. Our results provide evidence that motor module generalization across walking and reactive balance is reduced post-stroke and associated with lower walking performance. This is true even if the number and consistency of motor modules do not differ with respect to neurotypical individuals, as shown in prior results. Moreover, as reactive balance is a brainstem mediated behavior, a lower number of common motor modules across walking and reactive balance is consistent with less automatic control of gait after stroke.

Accumulating evidence suggests that motor module generalization may help to distinguish important and clinically-relevant differences in walking performance across individuals. Prior studies have demonstrated that various motor module-based metrics can describe differences in walking performance, such as motor module number (Clark et al. 2010; Fox et al. 2013; Rodriguez et al. 2013; Steele et al. 2015), consistency (Sawers et al. 2015; Allen et al. 2017), and recruitment timing (Routson et al. 2013). However, we did not identify differences in these metrics in our sample of stroke survivors. Instead, we found that motor module generalization across walking and reactive balance was reduced (**Fig. 2**). Motor module generalization, defined as recruiting a common set of motor modules across different movement behaviors, has previously been examined across different locomotor behaviors to understand limitations in the control of walking post-stroke (Routson et al. 2014). Although maintenance of balance is critical for walking, no study has examined generalization across walking and balance in stroke survivors with walking impairments. In two recent studies, we provided evidence that the amount of motor module generalization across walking and reactive balance is associated with walking performance. In particular, improved walking performance is accompanied by increased motor modules generalization after long-term training in healthy young adults (Sawers et al. 2015) and rehabilitation in individuals with Parkinson’s disease (Allen et al. 2017). Here, we show that higher levels of motor module generalization across walking and balance are also associated with better walking performance in stroke survivors. Taken together, these studies add to our understanding of how walking is controlled, providing compelling evidence that motor module generalization across walking and balance underlies well-coordinated walking. This suggests that utilizing a generalizable control strategy enables better walking performance.

Generalization of motor modules across walking and reactive balance may reflect automatic control of walking and be beneficial for robustly responding to external disturbances during walking. Rapid changes in the coordination of muscles is required to recover from discrete perturbations, such as those experienced by participants in the current study, and are thought to be governed by brainstem circuits (Stapley and Drew 2009). We previously demonstrated in healthy young adults that a common set of motor modules are recruited across walking and reactive balance (Chvatal and Ting 2012, 2013), suggesting a convergence of control on the automatic brain-stem mediated motor modules. Having a stroke, depending on the location of the infarct and affected brain areas, may disrupt the neural pathways governing the control of walking such that they no longer converge on the automatic brain-stem mediated motor modules. A loss of the ability to recruit the automatic reactive balance motor modules during walking post-stroke is consistent with reduced gait automaticity in this population (e.g., using dual-task paradigms, Hyndman et al. 2006; Liu-Ambrose et al. 2007; Plummer-D’Amato et al. 2008; Dennis et al. 2009). Less automatic control of walking is typically associated with increased stride-to-stride variability (Springer et al. 2006), perhaps due to neural commands that are less appropriate and more variable. Indeed, we found that the structure of the behavior-specific motor modules was more variable and less distinct than of the modules common to both walking and reactive balance (**Fig. 3**). Future study is needed to investigate how stroke location affects motor module generalization across walking and reactive balance and its relationship to gait automaticity.

The specific motor modules that are generalized across walking and reactive balance may further explain deficits in walking performance post-stroke. We identified three motor modules that were frequently generalized across walking and reactive balance (**Fig. 4**) that resemble motor modules previously identified during walking in healthy adults (Clark et al. 2010): an independent plantarflexor module, a proximal extensor module, and an ankle dorsiflexor module. Recruiting an independent plantarflexor motor module in the paretic leg across both walking and reactive balance was identified in our study as a significant predictor of walking speed post-stroke. That this module is important for walking speed is consistent with the role of the plantarflexors in generating forward propulsion (Liu et al. 2008; Neptune et al. 2008) and for successful walking performance post-stroke (Routson et al. 2013). The other motor modules frequently generalized across walking and reactive balance may be important for other measures of walking performance, such as better swing leg control when recruiting the dorsiflexor module.

Examining muscle recruitment across movement behaviors with different biomechanics and neural control may reveal important insights into changes in neuromuscular control underlying motor performance that might otherwise be overlooked. Although prior studies have examined the similarity of motor modules recruited across locomotor behaviors (Ivanenko et al. 2005; Fox et al. 2013; Routson et al. 2014), the similarity in motor modules may be heavily influenced by the similarity in biomechanics. In contrast, that there are common modules recruited for walking and standing reactive balance despite their different biomechanics and neural control circuitry provides strong evidence for an underlying neural strategy of generalization of muscle coordination patterns. This strategy appears to be compromised in stroke survivors and related to impairments in walking performance. Moreover, generalization across walking and reactive balance was not only reduced the in the paretic leg of stroke survivors but also their nonparetic leg (**Fig. 2**). Reduced generalization in the nonparetic leg may be due to either a neural deficit and/or a compensatory strategy to overcome deficits in the paretic leg and would not have been identified by examining walking alone. Indeed, examining movement behaviors that may utilize different biomechanics and neural pathways may be useful for understanding whether recruitment of muscles remains intact. Several of our stroke survivors exhibited classic patterns of motor module merging during walking, such as the merging of the ankle plantarflexors with more proximal hip and knee extensors (Clark et al. 2010). In some of these individuals (e.g., **Fig. 5**), the ankle plantarflexors were independently recruited during reactive balance despite their merged pattern in walking. Although this phenomenon could simply be an artifact of the methodology used to select the number of motor modules (four in balance, three in walking), we can generally rule this out because the plantarflexors were still merged with the proximal hip and knee extensors when four motor modules were extracted from walking and the independent plantarflexor module during reactive balance remained even when the number of motor modules extracted was reduced to three. Whether the merged plantarflexor control during walking was due to a choice (i.e. developed compensatory strategy) or a constraint (i.e., altered neural pathway integrity during walking) is unclear. Nevertheless, these results suggest that this individual retains the capability to independently recruit the plantarflexor in some capacity and might be more likely to regain independently plantarflexor control during walking through rehabilitation. This example, as well as main results showing changes in generalization across walking and reactive balance, demonstrate the information that can be gained about neuromuscular impairments limiting walking performance by examining multiple movement behaviors.

**Figure 5:**
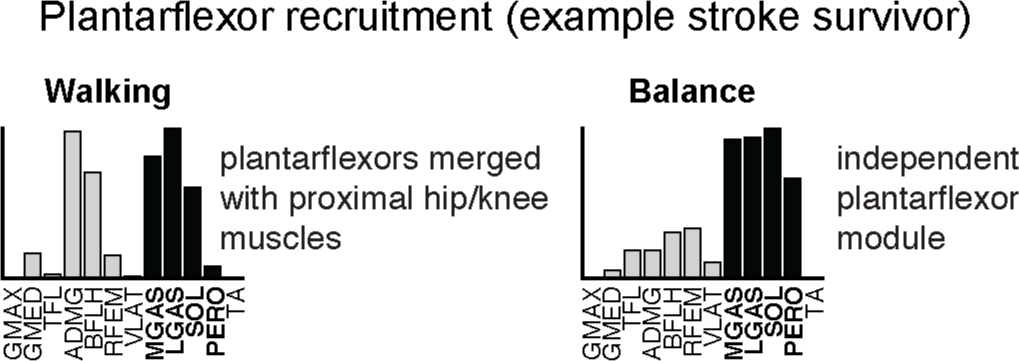
Example stroke survivor illustrating merged plantarflexor module during walking but independent recruitment of the plantarflexors in reactive balance.

